# Changes in group size during resource shifts reveal drivers of sociality across the tree of life

**DOI:** 10.1101/2020.03.17.994343

**Authors:** Albert B. Kao, Amanda K. Hund, Fernando P. Santos, Jean-Gabriel Young, Deepak Bhat, Joshua Garland, Rebekah A. Oomen, Helen F. McCreery

**Author notes:** corresponding authors: ABK, HFM.

## Abstract

From biofilms to whale pods, organisms have repeatedly converged on sociality as a strategy to improve individual fitness. Yet, it remains challenging to identify the most important drivers—and by extension, the evolutionary mechanisms—of sociality for particular species. Here, we present a conceptual framework, literature review, and model demonstrating that the direction and magnitude of the response of group size to sudden resource shifts provides a strong indication of the underlying drivers of sociality. We catalog six functionally distinct mechanisms related to the acquisition of resources, and we model these mechanisms’ effects on the survival of individuals foraging in groups. We find that whether, and to what degree, optimal group size increases, decreases, or remains constant when resource abundance declines depends strongly on the dominant mechanism. Existing empirical data support our model predictions, and we demonstrate how our framework can be used to predict the dominant social benefit for particular species. Together, our framework and results show that a single easily measurable characteristic, namely, group size under different resource abundances, can illuminate the potential drivers of sociality across the tree of life.

## Introduction

Living in a group can benefit an organism in numerous and diverse ways^1–4^. Sociality may improve an organism’s caloric intake^5–8^, reduce the costs of maintaining homeostasis^9–11^, decrease predation risk^12–15^, facilitate the discovery or defense of higher quality habitats or breeding sites^16–18^, increase the probability of finding a mate or opportunities for extra-pair copulations^4, 19, 20^, or increase the survival probability of offspring^18, 21, 22^. However, there are also many potential costs of group living, including increased competition over resources, disease risk^4^, and probability of infanticide^23, 24^.

A large body of literature explores how particular costs and benefits may affect group size and other social metrics^25–30^, yet it remains challenging to identify the most important drivers of sociality in particular species. For a given species, a subset of the potential benefits and costs will tend to dominate, and the balance of their magnitudes will determine the overall selection pressure for an organism to be social or asocial (as well as the typical group size)^31–33^. Knowledge of the underlying drivers of sociality is crucial not only for understanding the present distribution of social species and group sizes across taxa, but also for understanding the major evolutionary pathways by which sociality has arisen^34^ (*i.e.*, whether sociality has generally been driven by a subset of potential benefits, with secondary benefits accruing subsequently), as well as the future of social species in the face of emerging natural and anthropogenic threats^35–39^.

In practice, measuring the selection pressures driving sociality is difficult. It may require performing one or more experiments for each potential benefit of sociality, and the magnitude of a benefit may vary nonlinearly with group size and across different environmental contexts^22, 25, 27, 29, 40–42^. As a result, it is likely infeasible to comprehensively measure all of the selection pressures governing sociality for any species. Instead, studies typically isolate and study only one of the proximate mechanisms at a time. Studying a mechanism in isolation may allow researchers to determine whether that mechanism offers a measurable benefit (or cost) to group living, but cannot provide insight into its magnitude relative to other potential benefits and costs. Therefore, this strategy is unlikely to reveal the dominant drivers of sociality.

Here, we demonstrate a new approach to the problem of determining the drivers of sociality, using literature reviews, a new conceptual framework, and stochastic and analytic models. In nature, when the abundance of resources (*e.g.*, food) shifts over relatively short time scales^43^, the mean group size has been empirically observed to shrink for some species^44^, increase for others^45^, and remain constant for yet other species. We hypothesize that these differences in the direction in which group sizes shift when resources become scarce may be due to differences in the dominant driver of sociality across those species^46, 47^. If true, then observing changes in group size when resource abundance shifts could be a relatively simple method to generate a shortlist of likely drivers of sociality for a given species. Further experiments could then target those specific candidate drivers, in order to identify the dominant driver of sociality for particular species, and across taxa. Because resource availability is relatively easy to alter in the lab and group size easy to measure, this may be an experimentally tractable strategy to rapidly identify the dominant drivers of sociality.

## Results

### The fundamental resource-related benefits and costs of sociality

We first surveyed the literature and compiled a list of potential benefits of sociality, focusing on those related to gaining and consuming resources. We took a broad definition of “sociality” to include any group-living organism in a wide range of taxa, including bacteria, protists, fungi, arthropods, fish, birds, and mammals. Despite myriad resource-related benefits described across taxa in the literature^48^, we found that they could be condensed into a small set of functionally distinct classes. In total, we identified six fundamental benefits of sociality related to gaining or using resources (Table 1). We also recorded a wide range of non-resource-related benefits and costs of sociality, but we collapsed them into a single class, despite their variety, because they are generally indistinguishable from one another when studying how group size changes under shifting resource abundance.

**Table 1.**
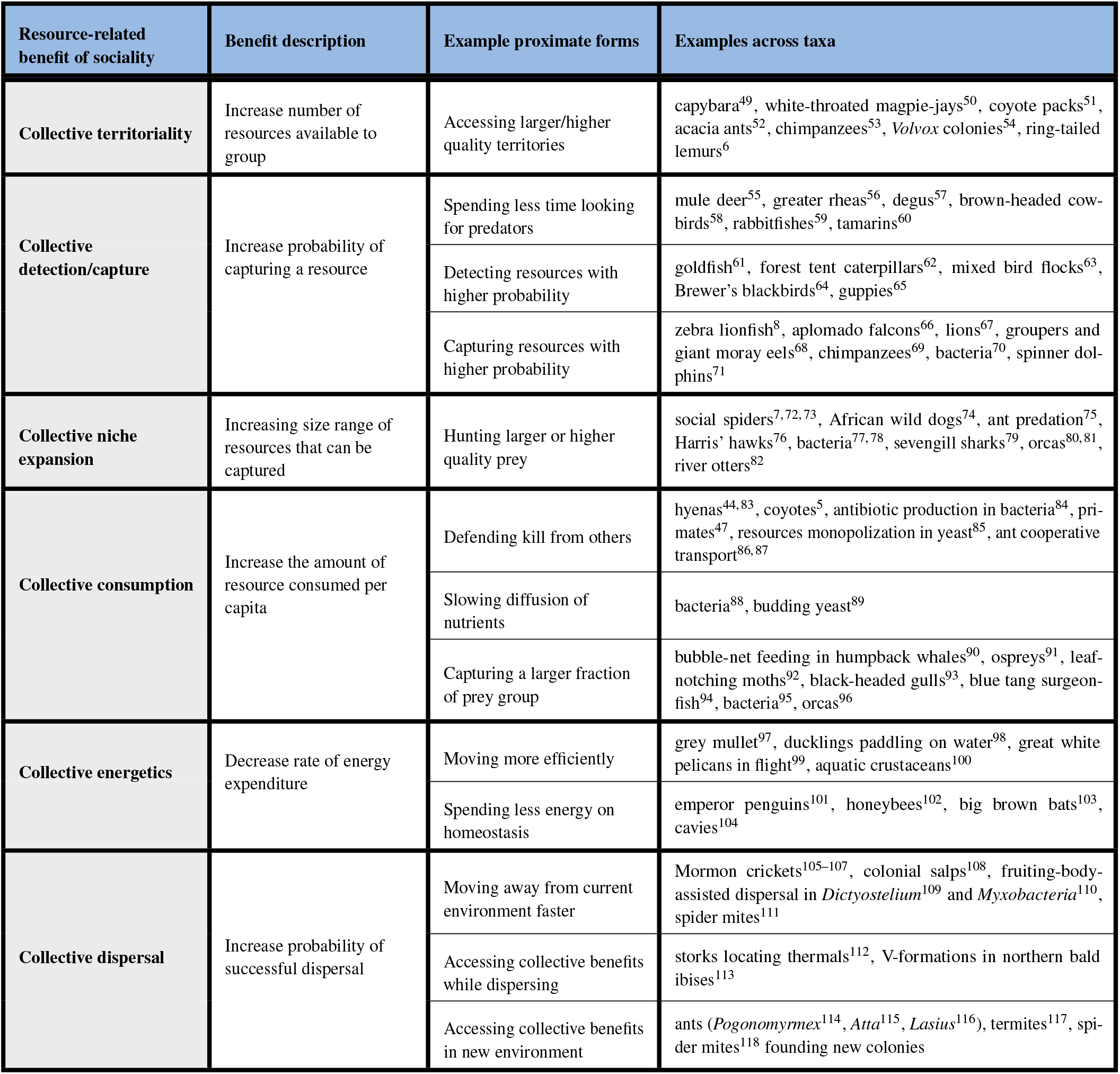
Six fundamental resource-related benefits of sociality. For each benefit of sociality, we give a brief description, note specific ways in which the benefit may manifest in different taxa, and list several examples of the benefit across diverse taxa. In addition to these six resource-related benefits, in our model we include the net effect of all non-resource-related benefits of sociality as a single benefit.

### Fundamental resource-related benefits of sociality

#### Collective territoriality

Larger groups are able to secure access to more resources. This describes organisms that can occupy larger territories, or those that can travel further as a group in order to explore a larger area. For example, larger groups of capybaras have larger and higher-quality home ranges^49^, while larger *Volvox* colonies can move farther in the water column to exploit a larger productive area^54^.

#### Collective detection/capture

Larger groups detect and capture a resource with higher probability. This benefit may occur because larger groups have a higher probability of detecting a resource (*e.g.*, individuals in larger groups spend less time looking for predators due to “shared vigilance,” allowing them to devote more time to foraging) or of capturing a detected resource. For example, degus spend less time individually on vigilance and more time foraging when in groups^57^, goldfish in larger shoals spend less time foraging before discovering a prey item^61^, and *Myxococcus* bacteria more efficiently predate on cyanobacteria at higher densities^70^.

#### Collective niche expansion

Larger groups can successfully capture larger or higher-quality resources. This benefit expands the dietary niche of the social species by giving members of large groups access to novel resources, rather than simply increasing the probability of capturing a resource that is also accessible to smaller groups (collective sensing/capture). This includes social spiders that use collective web structures to capture larger prey^7, 72, 73^, and some pathogenic bacteria that suppress their virulence until a quorum is reached, in order to overcome a host’s immune responses^77, 78^.

#### Collective consumption

Individuals in larger groups can consume more, per capita, of a captured resource before it is lost. Resources may be lost by being stolen by conspecifics or heterospecifics, or by environmental forces. Larger groups, therefore, may be able to better defend resources from others, as with coyotes^5^, or ants that utilize cooperative transport, carrying large resources to the nest, thus removing them from competition^86, 87^. Alternatively, collective feeding of a resource comprised of many components can permit the consumption of a larger fraction of that resource, such as biofilms that encapsulate a resource and slow the diffusion of nutrients^88, 89^, or humpback whales that use bubble nets to feed on a school of fish^90^.

#### Collective energetics

Individuals in larger groups expend less energy, on average, per unit time. This benefit may be due to aerodynamic or hydrodynamic benefits, such as with great white pelicans, which have reduced energy expenditure while flying in a V-formation^99^. Other social animals are better able to thermoregulate in a large group, such as emperor penguins^101^ and honeybees^102^, lowering the energetic costs of maintaining homeostasis.

#### Collective dispersal

Larger groups can more easily disperse from a poor habitat to a new habitat. This benefit may arise because groups move more efficiently while searching for a new environment, such as Mormon crickets that move more ballistically as a swarm^105–107^ and slime molds that self-assemble into a dispersal stalk^109^. Alternatively, larger groups may disperse more successfully by accessing one or more of the other collective benefits when traveling (such as storks saving energy by collectively sensing air thermals^112^) or when they arrive at their new habitat (such as some ant species that exhibit increased survival of new colonies when multiple queens found nests together^114–116^).

### Fundamental resource-related cost of sociality

#### Intragroup competition

Individuals in larger groups suffer increased competition for resources. While the details of intragroup competition differ among species (*e.g.*, the size of the finder’s share, whether dominant individuals claim a larger share, whether group mates are subject to scramble or contest competition), in general there will be fewer resources available per capita as group size increases.

### Non-resource-related benefits and costs of sociality

In addition to these fundamental resource-related benefits and costs of sociality, there are a large number of other effects of sociality that are unrelated to resource acquisition. Such benefits include avoiding predation by detecting predators more reliably (the “many eyes” hypothesis), evading predators more effectively (“Trafalgar” or “confusion” effects), shunting risk onto groupmates (the “selfish herd” principle), and diluting the overall risk of predation^4^. Sociality may also affect the probability of acquiring mates (such as in leks^119^, and harems^120^), increase fitness via cooperative breeding^121, 122^, decrease fitness via inbreeding^123^, and alter parasite risk^124–128^. In short, there are a large number of potential benefits and costs of sociality that are not directly related to resource acquisition, which we combine into a single class (<u>*other mortality*</u>).

All of the sociality mechanisms described here, resource-related or not, may in principle result in net benefits or costs for some species and contexts. For example, the probability of capturing prey may shrink as group size increases if the prey can more easily detect a larger group. For simplicity, we assume that competition represents a cost of sociality and, for all other mechanisms, we focus on the net benefits of sociality.

### Modeling how benefits of sociality and resource abundance affect group sizes

With this understanding of the fundamental benefits of sociality, we can investigate how these benefits may impact group size when resource abundance shifts. To do so, we introduce a general model of a group foraging for resources, which we briefly describe qualitatively here, depict schematically in Figure 1, and describe quantitatively in *Methods: Detailed description of the model framework*.

**Figure 1.**
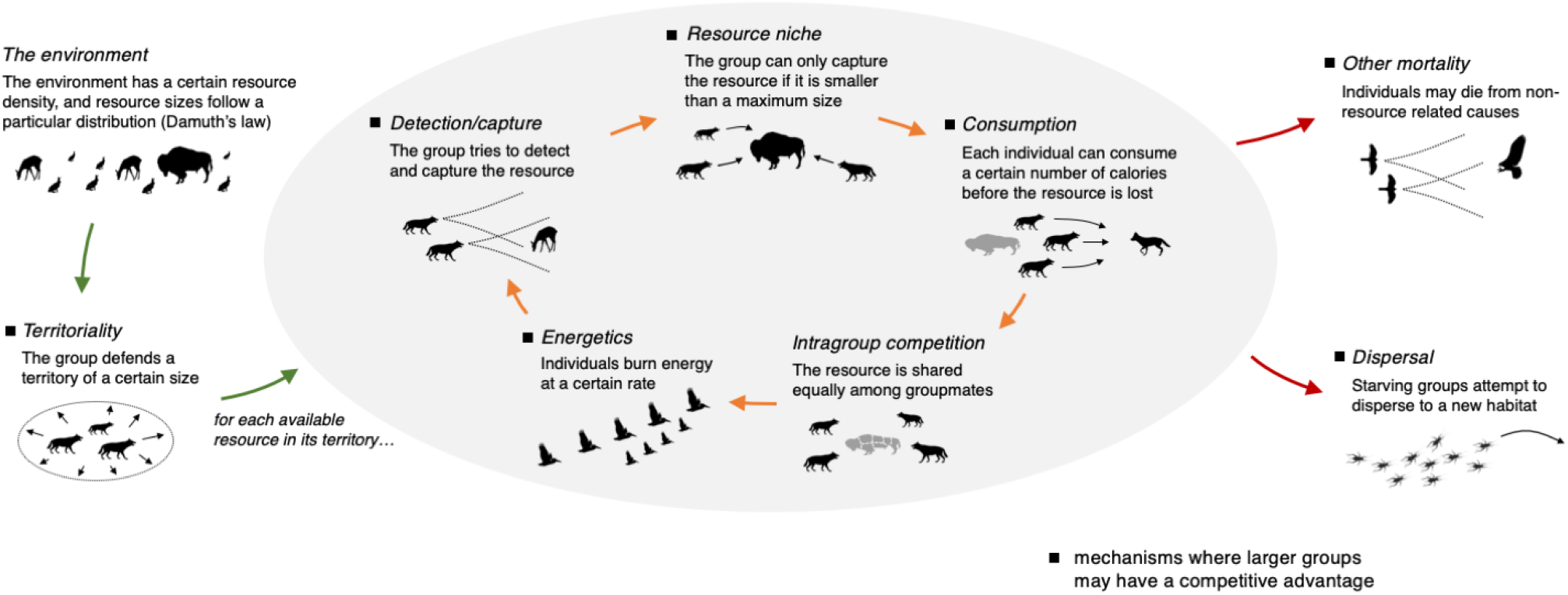
Modeling framework for investigating the effects of the benefits of sociality and resource abundance on group sizes. We specify an environment, which consists of a particular resource density and a power-law resource mass-abundance relationship. A group is allocated a territory of a certain size, which provides it access to a certain number of potential resources. The group then tries to detect, capture, and consume each resource in its territory, spending energy in the process. If the individuals in the group run out of energy, then the group attempts to disperse to a new habitat. During a simulation, individuals may also die from non-resource-related causes. The probability of surviving a simulation, as a function of group size, the dominant benefit of sociality, and resource abundance, is recorded. Although mammals or birds are typically used here as an illustration of different parts of the model framework, we stress that this framework can be used to model a wide variety of taxa.

In the model, a group exists in an environment containing resources whose sizes follow a power-law distribution, based on empirical and theoretical work on mass-abundance relationships in nature^129–131^ (Figure 1, *the environment*). Consequently, during a simulation a group typically has access to many small resources and few large resources. A group is assigned a territory of a certain size, giving it access to a particular number of potential resources (Figure 1, *territoriality*).

At the start of the simulation, each individual in the group has a store of energy. During the simulation, the group attempts to detect, capture, and consume each of the potential resources. There is some probability that the group successfully detects and captures a particular resource (Figure 1, *detection/capture*); however, the group is limited to those resources that are within its niche (Figure 1, *resource niche*). If a resource has been successfully captured, each individual in the group extracts some energy from it, based on how much can be consumed before it is lost (Figure 1, *consumption*), and assuming that individuals in the group share resources equally (Figure 1, *intragroup competition*). As the group tries to capture and consume the available resources, each individual burns a certain amount of energy (Figure 1, *energetics*), and there is some probability that an individual dies due to a non-resource-related cause (Figure 1, *other mortality*). If the individuals in the group run out of energy during the simulation, then the group attempts to disperse to a new habitat; it successfully disperses with some probability and otherwise the individuals in the group die (Figure 1, *dispersal*). Therefore, an individual survives a simulation if 1) it does not die from a non-resource-related cause and does not run out of energy, or 2) it does not die from a non-resource-related cause and runs out of energy but successfully disperses.

For each simulation, we “turned on” one of the potential benefits of sociality, such that larger groups exhibit improved performance in that aspect of the simulation (Figure 1, black squares). We ran simulations at different group sizes, and across a wide range of all of the free parameters of the model, turning on each of the benefits individually (for simplicity and interpretability, we did not consider multiple potential benefits acting in concert, though our model could be used to perform such analyses in the future). For each combination of parameter values, we ran 10,000 simulations for each group size ranging from 1 to 100 and computed the optimal group size that maximized survival probability, for both an “abundant resources” condition and a “scarce resources” condition (*Methods: Computing the optimal group size*).

### The effect of the sociality benefit class on group size shifts

Depending on which benefit of sociality was present in the simulation, we found that the optimal group size can decrease, increase, or stay the same when resources become scarce. In Figure 2A, we show illustrative examples of the optimal group size decreasing (when the collective detection/capture benefit of sociality was present) and increasing (when the collective consumption benefit of sociality was present). In addition to our stochastic simulations, we also developed an analytical solution to our model (exact when the number of resources *μ* = 1, see *Methods: Analytical solution*), which closely matches our simulation results (Figure 2A).

**Figure 2.**
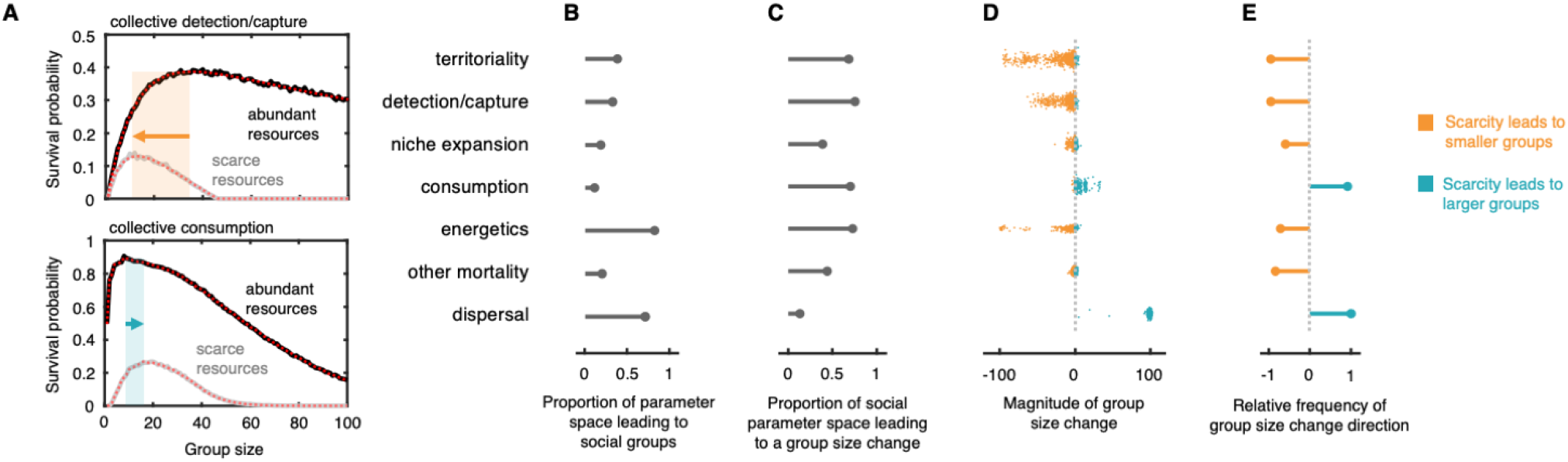
The benefits of sociality can cause either increases or decreases in group size when resources become scarce. **A.** Resource scarcity can lead to decreases (top) or increases (bottom) in the optimal group size, depending on the dominant benefit of sociality and the parameters of the environment. In the top panel, collective detection/capture is the dominant benefit, with parameters *μ* = 1, 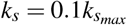, *s* = 1.01, *α*_*s*_ = 0.01, *α*_*h*_ = 0.5, and *α*_*d*_ = 0.999, while in the bottom panel, collective consumption is the dominant benefit, with parameters *μ* = 20, *γ*_*d*_ = 0.5*γ*_*max*_, *s* = 1.01, *α*_*s*_ = 0.5, *α*_*h*_ = 0.25, and *α*_*d*_ = 0.001 (see *Methods: Detailed description of the model framework* for parameter definitions). Black and grey lines show results from simulations (mean of 10,000 repetitions) for the abundant and scarce resource conditions, respectively, while the red and pink lines show the analytical solution for the same parameter values as in the simulations. **B.** The proportion of parameter space where the optimal group size was greater than 1 for either the abundant or the scarce resource condition, for each benefit of sociality. **C.** The proportion of social parameter space (i.e., the fraction of parameter space shown in B) that leads to a change in group size. **D.** Scatter plot of the resulting group size shift for those sets of parameter values that led to a group size shift (orange points indicate that group size decreases when resources are scarce, blue points indicate that group size increases when resources are scarce; a small amount of jitter was added in both the horizontal and vertical directions to more clearly visualize the data points). **E.** The relative frequencies of the shift direction for those sets of parameter values that led to a group size shift (a value of −1 indicates that all group size shifts were decreases in group size when resources are scarce, +1 indicates that all group size shifts were increases in group size when resources are scarce).

Across all of parameter space, a given benefit of sociality led to social groups (*i.e.*, survival probability was maximized when *M* > 1, for either the abundant or scarce resource condition), in 11 to 82% of sets of parameter values, depending on the active social benefit (Figure 2B). In subsequent analyses, we focused on those sets of parameter values that permitted social groups, since our aim was to study social, rather than asocial, species.

Within this restricted region of parameter space, between 13 to 76% of sets of parameter values led to a shift in group size when resource abundance shifted (Figure 2C), with the territoriality, detection/capture, consumption, and energetics benefits most frequently leading to shifts in group size. Furthermore, the magnitude and direction of group size shifts strongly depended on which benefit of sociality was present (Figure 2D). The territoriality and detection/capture mechanisms tended to lead to decreases in group size when resources became scarce, with a wide range of magnitudes, depending on the parameter values. By contrast, the energetics mechanism also generally resulted in decreases in group size under scarcity, but the magnitudes tended to be either small or large. While the niche expansion and other mortality mechanisms could lead to either decreases or increases in group size, the magnitudes of these shifts tended to be small (and therefore possibly difficult to detect in experiments). The consumption and dispersal mechanisms tended to lead to group size increases under scarcity, with collective dispersal leading to very large increases in group size (however, for this mechanism a wide range of group sizes had nearly identical survival rates, so group size changes in nature or in experiments may be smaller than shown in Figure 2D; see *Methods: Computing the optimal group size*). We further quantify group size shifts under scarcity by plotting the relative frequency of decreases and increases in group size (Figure 2E), confirming that the consumption and dispersal benefits are the only two that robustly cause increases in group size under scarcity, while all of the other benefits tend to lead to decreases in group size (though with widely varying magnitudes).

### Empirical evidence for the model predictions

We returned to the literature to examine the extent to which existing data support, or contradict, our theoretical predictions. Based on the quantitative results in Figure 2, we partitioned the seven benefits of sociality into three groups: those that tend to lead to decreases (territoriality, detection/capture, and energetics), increases (consumption and dispersal), or only very small changes in group sizes (niche expansion and other mortality) when resources become scarce (Figure 3, “Model results”). The fact that the seven benefits of sociality evenly partition into these three groups suggests that a single measurement (direction of group size shift under scarcity) can be a highly informative indicator of the dominant driver of sociality for a species by efficiently selecting a subset of the possible benefits. We then looked for empirical evidence of group size shifts when resource abundance shifts.

**Figure 3.**
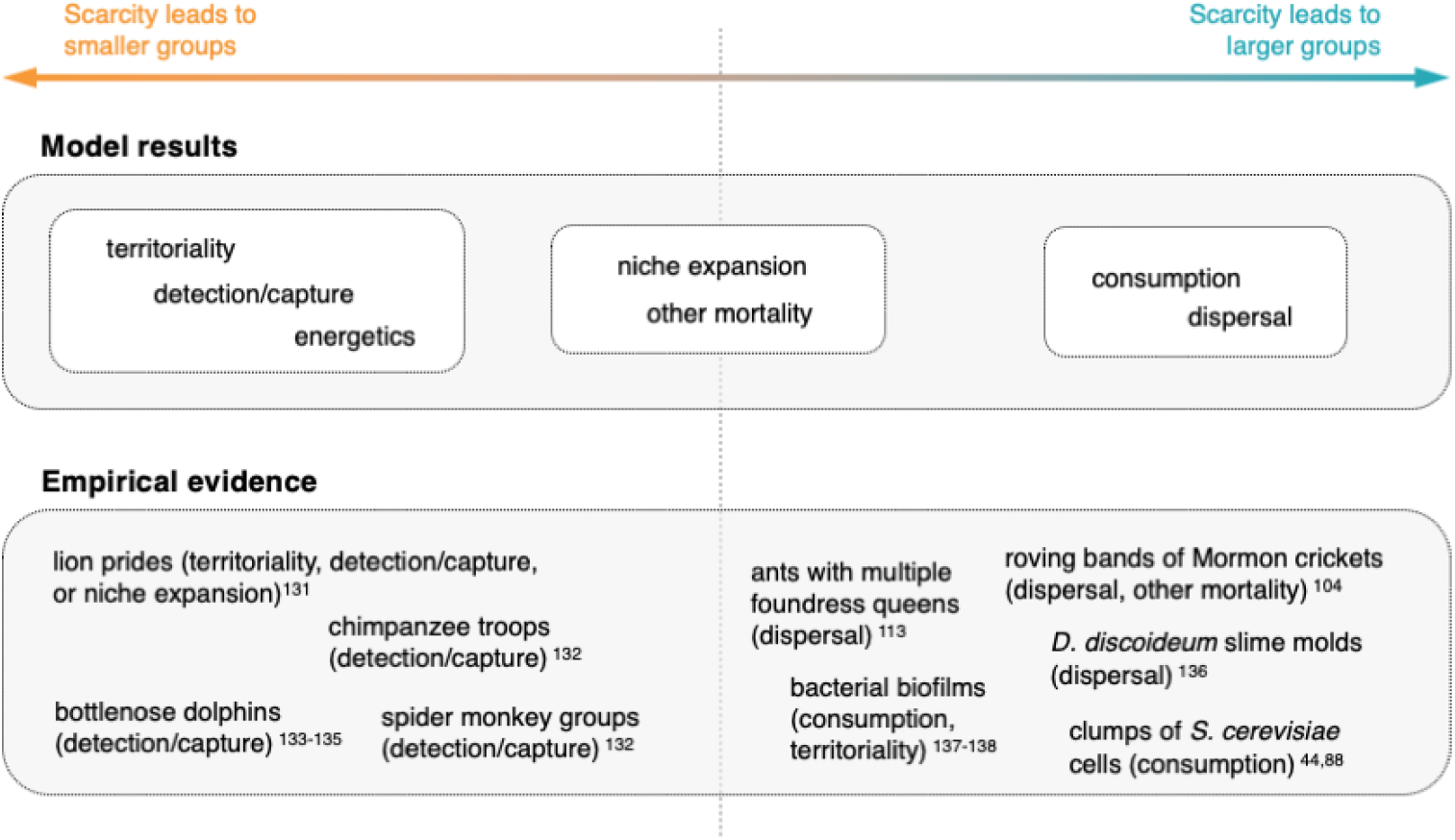
Comparing our model predictions to existing empirical data. The model results, shown in Figure 2B-E, lead to broad classifications of the benefits of sociality based on their general tendency to produce either decreases, increases, or only minor shifts in group size when resources become scarce. Existing empirical evidence, while relatively scant, tend to agree with our model predictions. We note that this comparison is preliminary, since even for the species that we list in this figure, the dominant benefit of sociality is not known with certainty.

While existing data are relatively scant, we found a high degree of agreement between empirical examples and our model predictions (Figure 3, “Empirical data”). For example, there is evidence that lion prides^132^, chimpanzee troops^133^, bottlenose dolphin pods^134–136^, and spider monkey troops^133^ are smaller when resources are scarce. The most plausible benefit of sociality for these species, as previously established from empirical data, is that group living improves the ability of individuals to capture prey or other resources, whether due to improved detection or capture of resources, increased territory size, or ability to capture larger resources.

By contrast, several species are known to increase their group size under scarcity. Large bands of Mormon crickets likely have an improved ability to discover new habitats (dispersal), but may also benefit from improved predator avoidance (other mortality)^105^. Slime molds, such as *Dictyostelium discoideum*, switch from a solitary to a social phase when food is scarce, dispersing via spores within an aggregate fruiting body (dispersal)^137^. For some species of ants, new colonies comprising multiple foundress queens are more likely to survive than single queens (dispersal)^114^. Bacteria can form biofilms^138, 139^ and the yeast *Saccharomyces cerevisiae* can form clumps^45, 89^ when nutrients are scarce, which absorb more of a resource’s nutrients and/or cooperatively produce digestive enzymes (consumption). Some of the species likely benefiting from collective dispersal exhibit the extremely large increases in group size that we observed in our model, such as Mormon crickets and locusts that transition from solitary individuals to vast swarms.

For other species, group size shifts when resource abundance shifts but with less clear evidence of the driver of sociality. For example, finches and monk parakeets form larger flocks when resources are scarce^140, 141^. These groups may benefit from improved information gathering and sharing (collective sensing/capture), but alternatively large groups may form simply due to aggregation at a smaller number of available food patches. Similarly, many gram-negative bacteria form biofilms only in nutrient-rich media and detach when nutrients become scarce, although the function of the biofilm is not known with certainty^142^. Hyenas were shown to decrease their group size under scarcity, with hypothesized mechanisms driving sociality including collective capture, collective consumption, and improved infant safety^44^.

While existing empirical data generally support our model predictions, we stress that this comparison is preliminary, as the dominant driver of sociality is not known with certainty for any species. For many species, there are multiple social mechanisms hypothesized to significantly affect individual fitness. In Figure 3, we listed species for which the set of possible mechanisms is reasonably small, or for which there is greater confidence among researchers about which is the dominant driver of sociality. We did not list in Figure 3 species with evidence of group size shifts under scarcity but with less known about the underlying drivers.

Experiments specifically designed to test our model predictions are necessary. A simple such test would be to change resource availability while keeping other aspects of the environment constant (such as patchiness), recording any changes in group size. Our model can produce a reduced subset of potential drivers of sociality based on the group size shift, and these potential drivers can be individually tested to measure the associated selection pressure. Our model predicts that at least one of these potential drivers will exhibit a substantially stronger effect compared to the other possible drivers of sociality.

### Generating predictions for particular species

In the analyses above, we aggregated the results of all of our simulations, which were conducted across a wide range of parameter values (see *Methods: Detailed description of the model framework*). We deliberately selected a wide range in order to capture the lifestyles of diverse organisms and environments across the tree of life. However, researchers wanting to identify the dominant driver of sociality for a particular species will likely have some knowledge of the specific region of parameter space that it inhabits. If so, then our model could be run within that specific region of parameter space and more targeted predictions could be generated for that species.

Here, we demonstrate how our model framework could be used to create a shortlist of likely drivers of sociality for particular species. To avoid making unsubstantiated assumptions about any real biological species, we showcase how to apply our framework to two mythological creatures: dragons and unicorns (Figure 4A,D). We assume that dragons are carnivorous, large, fierce, can fly, and can breathe fire. By contrast, unicorns are herbivorous, smaller, timid, live in forests, and we assume that their horns result mainly from sexual selection and are not effective defensive tools. Based on these traits, we identified hypothetical parameter ranges for both species, as a researcher would do for their focal real species-see *Methods: Dragon and unicorn analyses* for a complete explanation of our chosen parameters for these examples.

**Figure 4.**
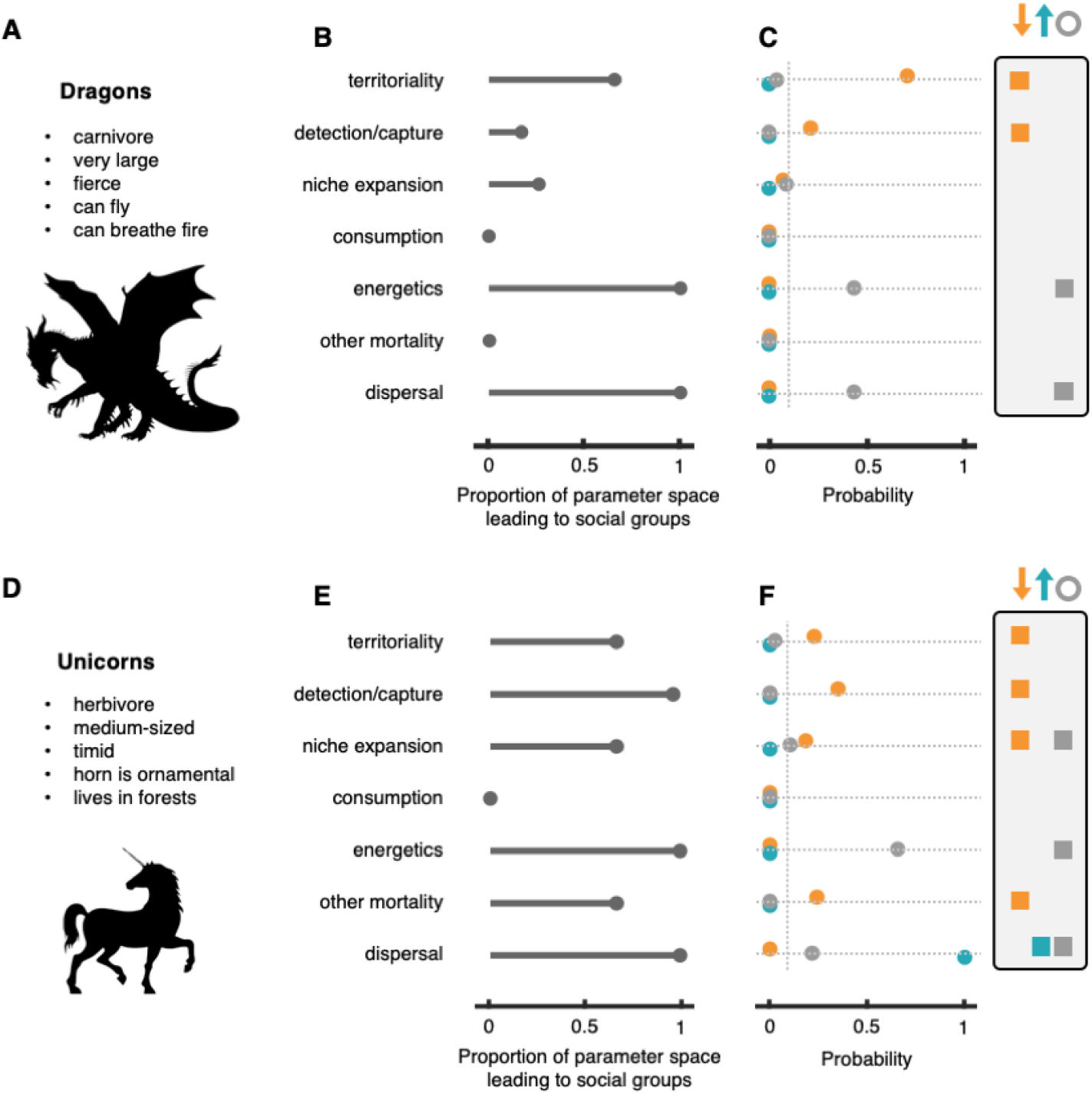
Model predictions may be taxon-dependent. We apply our model to two mythological creatures, dragons and unicorns. **A.** Dragons are large, can fly, and can breathe fire. **B.** The proportion of dragon parameter space where the optimal group size was greater than 1 for either the abundant or the scarce resource condition, for each benefit of sociality. **C.** The probability that observed group size changes arise from each benefit type. The x-axis shows the proportion of scarcity-caused group size decreases (in orange), increases (in blue), and simulations with no change (in grey) attributable to each benefit type, for the dragon parameter space. Inset at right highlight benefits that surpass 0.1 probability of driving a given result. Parameters are described in *Methods: Detailed description of the model framework*, and the dragon parameter space for these analyses was estimated as described in *Methods: Dragon and unicorn analysis*. **D.** Unicorns are medium-sized, timid, and have only ornamental horns. **E.** The proportion of unicorn parameter space where the optimal group size was greater than 1 for either the abundant or the scarce resource condition. **F.** The probability that observed group size changes arise from each benefit type for the unicorn parameter space.

We can narrow down the list of likely drivers of sociality for dragons and unicorns even without collecting any additional data, simply by knowing that these two species are social and reside in particular regions of parameter space. Within the dragon region of parameter space, our model predicts that collective consumption and other mortality benefits can never lead to social groups (optimal group size > 1) regardless of resource abundance (Figure 4B). Since we presume that dragons can be found in groups (because we are asking why dragons are social), we can eliminate these benefits as likely sociality drivers. For unicorns, on the other hand, we can eliminate only collective consumption with this logic (Figure 4E).

To narrow the list of potential drivers even further, we took a likelihood approach. Supposing a researcher observes how group size changes when resources become scarce, we can assign a probability that each of the potential benefits of sociality resulted in the observed outcome. In other words, if we observe a decrease (or increase or no change) in dragon group size when resources become scarce, what proportion of simulations resulting in decreases were caused by each potential benefit? Considering only the easily-measurable direction, rather than the magnitude, of group size shifts, we highlight those mechanisms that had a probability greater than 0.1 of generating the given group size shift. In the case of a decrease in group size for dragons, for example, our model predicts that collective territoriality and collective detection/capture are the most likely drivers of sociality (orange dots and squares in Figure 4C). If no change in group size is measured, collective energetics and collective dispersal are the most likely drivers (grey dots and squares). The model predicts that increases in group size should not be observed in the region of parameter space that dragons reside, regardless of the driver of sociality (blue dots in Figure 4C). If unicorn groups decrease in group size under scarcity, our model highlights four potential drivers of sociality: collective territoriality, collective detection/capture, collective niche expansion, and other mortality (orange dots and squares in Figure 4F). However, consistent with Figure 2, for collective niche expansion and other mortality we expect only small decreases in group size, which may be more difficult to measure in practice. An increase in group size would suggest the collective dispersal mechanism as the dominant driver of sociality (blue dots and squares), while no change in group size highlights the collective niche expansion, collective energetics, and collective dispersal mechanisms (grey dots and squares in Figure 4F).

Overall, these results demonstrate how predictions can differ depending on the region of parameter space in which a species resides. Simply by knowing that the species under study is social, or additionally, knowing how group sizes shift when resources become scarce, one can dramatically reduce the list of likely drivers of sociality. Researchers could then test the remaining sociality drivers to identify the main drivers of sociality for that species.

## Discussion

We described here a novel conceptual framework—applicable across diverse taxa, from bacteria to large mammals—to study how a variety of potential benefits of sociality can affect group size (Figure 1). While there are a large number of potential benefits related to gaining resources, our analysis condensed these benefits into six classes with fundamentally different functional forms (Table 1). Stochastic simulations of our model of groups acquiring resources, and its analytic solution, revealed that the direction and magnitude of group size changes due to declines in resource abundance are strongly dependent on which benefit is the underlying driver of sociality (Figure 2). This allowed us to partition the benefits of sociality into three groups: those that tend to lead to smaller groups under scarcity (collective territoriality, collective detection/capture, and collective energetics), those that tend to lead to little or no change in group size (collective niche expansion, other mortality), and those that tend to lead to larger groups under scarcity (collective consumption, collective dispersal) (Figure 3). Although currently limited, existing data on a variety of species appear to broadly support our model predictions. Finally, we demonstrated how a researcher could apply our framework to particular species and how the model can make different predictions depending on the relevant parameter space for that species (Figure 4).

Our approach of taking a bird’s eye view of all resource-related benefits of sociality across taxa necessitated making simplifying assumptions. We implicitly assume that resources are randomly distributed in the environment and can only shift in abundance, that available resources follow a particular mass-abundance relationship, and that the abundance of all resources declines uniformly under scarce conditions. In real habitats, resources can also vary in patchiness, may follow different mass-abundance relationships, and the abundance of resources may decline nonuniformly (*e.g.*, large resources may be more or less resilient during droughts). Furthermore, resources may shift not only in abundance but also in frequency and predictability. We also do not model intragroup dynamics (*e.g.*, dominance hierarchies, finder’s shares, division of labor, etc.), intergroup competition, interspecies interactions, demographic structures, and other strategies besides dispersal to stave off starvation. In our model, we assumed that all of the mechanisms, except intragroup competition, function as potential benefits of sociality, rather than potential costs. Despite—and because of—these simplifications, our model highlights fundamental dynamics governing group size across species. Nonetheless, our model framework is general enough to accommodate these additional features if they are particularly important to a species of interest.

An important direction for future research is to apply our modeling framework to understand how multiple benefits of sociality may interact with each other. Many social species are likely to benefit from multiple mechanisms, such as collective territoriality, collective detection/capture, and collective niche expansion. These may combine nonlinearly to promote sociality and affect group size shifts and may alter our predictions of what group sizes we expect to observe in nature.

Importantly, our framework reveals that group size shifts under changes in resource availability can be an insightful clue about the underlying driver of sociality. Rather than performing separate experiments on each possible benefit of sociality (*i.e.*, the many proximate forms, some of which are listed in Table 1), we argue that simply observing group size changes can substantially reduce the set of likely drivers of sociality. This experiment is often tractable and cost-effective to perform in the lab. In this work, we reported changes in optimal group size, but some groups may have a relatively fixed group size, and may not adjust over the course of a typical experiment. In such cases, researchers could use proxies for optimal group size, such as levels of stress hormones in groups of different sizes^6^. Comparative phylogenetic approaches, such as correlating group size to typical habitat conditions, could also be fruitful for taxa with relatively fixed group sizes, for which we expect changes to occur on evolutionary timescales.

Knowing the major drivers of sociality across species is crucial to a comparative understanding of how sociality evolves across diverse biological systems^143^, as well as anticipating the resilience of social species when facing environmental change. For example, we observe a possible correlation between organism size and group size change, or likely sociality driver (Figure 3); future research on this correlation could explore its implications for the evolution of sociality and social resilience. Our model framework demonstrated that the different benefits of sociality differ in their propensity to generate social groups (Figure 2B). Improving our understanding of which benefits underlie sociality among extant species will allow us to test whether some benefits tend to drive the evolution of sociality, with other benefits accruing secondarily. In addition, this framework could be applied at the sub-organismal level, potentially explaining the group sizes of cellular aggregations such as sperm cells^144, 145^, erythrocytes and platelets^146–148^, and some immune cells^149–151^. Taking a high-level view of sociality across the tree of life, while abstracting away many of the finer details, may therefore be a powerful approach towards a deeper understanding of one of the major evolutionary transitions.

## Acknowledgements

ABK acknowledges support from a Baird Scholarship, and ABK and JG acknowledge support from an Omidyar Fellowship from the Santa Fe Institute. AKH, J-GY, FPS, RAO, and HFM acknowledge support from a James S. McDonnell Postdoctoral Fellowship Award. DB gratefully acknowledges financial support from NSF Grant No. DMR-1608211. The James S. McDonnell Foundation and the Santa Fe Institute additionally funded meetings and a working group grant for this work. Andrew Berdahl contributed ideas (particularly on the ballistic motion of locust swarms) that stimulated initial discussions for this project.

## Data availability

We will publish the data and code associated with our model upon publication.

## Author contributions

ABK conceived of the study. AKH, HFM, ABK, FPS, and RAO performed the literature review to identify the fundamental drivers of sociality. AKH, HFM, and RAO performed the literature review to compare the model predictions to existing empirical evidence. All authors contributed to the conceptual development of the model. ABK, FPS, JG, J-GY, and DB and wrote simulation code, and ABK ran the simulations and generated the figures. J-GY, DB, and FPS developed the analytical solution to the model. ABK, HFM, AKH, FPS, J-GY, and RAO drafted the manuscript. All authors edited the manuscript and gave final approval for publication.

## Methods

### Detailed description of the model framework

First, we specify the resource environment within which the group forages. The environment contains a certain density of resources; these resources vary in size from 0 to *C*_*max*_ = 1, and the probability density *P*(*C*) that a resource is a certain size *C* is given by 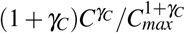, where we set the exponent to *γ*_C_ = −3/4, following existing empirical and theoretical work on mass-abundance relationships in nature, known as Damuth’s law^129–131^. A group is assigned a territory, the area of which is proportional to the number of potential resources, *μ*, that the group can access during the simulation.

Each individual in a group of size *M* begins with *c*_0_ energy. During the simulation, the group attempts to detect, capture, and consume each of the *μ* potential resources. There is some probability *p*_*s*_ that the group successfully detects and captures that resource. However, the group can capture a resource only if it is within its niche, *i.e.*, *C* ≤ *C*_*h*_, where *C*_*h*_ is the maximum size resource that the group can capture; the group must ignore the resource if it is too large. If the resource has been successfully captured, then each individual in the group consumes some portion of the resource. Each individual can potentially consume *c*_*d*_ of the resource before it is lost (*e.g.*, to conspecifics, heterospecifics, or the environment). However, because we assume that groupmates share resources equitably, individuals can each consume a maximum of *C*/*M* of the resource (*i.e.*, if the entire resource is consumed). Therefore, the actual amount of the captured resource that an individual consumes is *c*_*c*_ = *min*{*c*_*d*_,*C*/*M*}; if the resource was not detected and captured, or was outside of its niche, then *c*_*c*_ = 0 for that resource.

While the group tries to capture the available resources, each individual burns a total of *c*_*b*_ energy. In addition, there is some probability *p*_*p*_ that an individual dies due to some non-resource-related cause. If the individuals in the group run out of energy during the simulation, *i.e.*, *c*_0_ + Σ*c*_*c*_ −*c*_*b*_ < 0, then the group attempts to disperse to a new habitat; it successfully disperses with some probability *p*_*e*_ and otherwise the individuals in the group die.

Therefore, an individual survives a simulation if 1) it does not die from a non-resource-related cause and does not run out of energy, or 2) it does not die from a non-resource-related cause and runs out of energy but successfully disperses.

For a given simulation, we “turned on” one of the potential benefits of sociality; depending on which mechanism is turned on, the magnitude of one of *μ*, *p*_*s*_, *C*_*h*_, *c*_*d*_, *c*_*b*_, *p*_*p*_, or *p*_*e*_ will be a function of group size *M*. We used generic functions to model how these variables change with *M*:

#### Collective territoriality

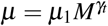, where *μ*_1_ is the number of potential resources available to a solitary individual, and *γ*_*t*_ is the strength of this social benefit.

#### Collective detection/capture

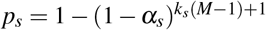, where *α*_*s*_ is the probability that a solitary individual detects and captures a resource within its niche, and *k*_*s*_ is the strength of this social benefit. This functional form, as well as similar ones below, are based on the “perfect many eyes” scenario^152^, but modified to allow for different strengths of the social benefit; this function increases from *p*_*s*_(*M* = 1) = *α*_*s*_ to *p*_*s*_(*M* = ∞) = 1.

#### Collective niche expansion

*C*_*h*_ = *α*_*h*_*C*_*max*_*M*^*γh*^, where *α*_*h*_ is the maximum size resource that a solitary individual can capture, as a proportion of *C*_*max*_, and *γ*_*h*_ is the strength of this social benefit.

#### Collective consumption

*c*_*d*_ = *α*_*d*_*C*_*max*_*M*^*γd*^, where *α*_*d*_ is the amount of a captured resource, as a proportion of *C*_*max*_, that a solitary individual can consume before the resource is lost, and *γ*_*d*_ is the strength of this social benefit.

#### Collective energetics

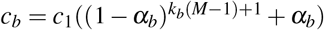 where *c*_1_ is the amount of energy that a solitary individual burns during the simulation, *k*_*b*_ is the strength of this social benefit, and *α*_*b*_ represents the fraction of energy burned by a solitary individual that an individual in an infinitely large group burns. This function decreases from *c*_*b*_(*M* = 1) = *c*_1_ to *c*_*b*_(*M* = ∞) = *α_b_c*_1_ (where 0 ≤ *α*_*b*_ ≤ 1).

#### Other mortality

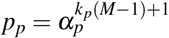, where *α*_*p*_ is the probability that a solitary individual dies from a non-resource-related cause, and *k*_*p*_ is the strength of this social benefit. This function decreases from *p*_*p*_(*M* = 1) = *α*_*p*_ to *p*_*p*_(*M* = ∞) = 0.

#### Collective dispersal

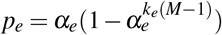, where *α*_*e*_ is the probability that an infinitely large group successfully disperses, and *k*_*e*_ is the strength of this social benefit. This function increases from *p*_*e*_(*M* = 1) = 0 to *p*_*e*_(*M* = ∞) = *α*_*e*_.

We set maximum values of the strengths of the social benefits, typically such that the benefit can only grow sublinearly with group size, with some exceptions (see *Methods: Limits on the strength of the social benefits*), and ran simulations for each mechanism, where the strength of the mechanism was set to 10%, 50%, or 90% of its maximum value. We set *μ*_1_ to either 1 or 20 to capture different ecological regimes (where, respectively, resources are relatively rare or common), and *α*_*s*_, *α*_*h*_, *α*_*d*_, *α*_*b*_, *α*_*p*_, and *α*_*e*_ to 0.001, 0.01, 0.1, 0.25, 0.5, 0.75, 0.9, 0.99, or 0.999. In general, the parameters *α*_*b*_, *α*_*p*_, and *α*_*e*_ affect dynamics only when their respective mechanisms are turned on, since *α*_*b*_ and *α*_*e*_ play a role only when those mechanisms are on, and we set *α*_*p*_ = 0 unless that mechanism is turned on.

We calculated the probability that an individual survives a simulation for group sizes ranging from 1 to 100, for both an “abundant resource” regime and a “scarce resource” regime. We define the abundant resource regime as the condition where a solitary individual is expected to have exactly 0 energy at the end of the simulation (see *Methods: Calculating the “edge of starvation”*). The free parameters available to tune this “edge of starvation” are *c*_0_ and *c*_1_. We choose to use *c*_1_, while setting *c*_0_ to 1. If the edge of starvation (and therefore the abundant resource regime) is given by 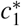, then we define the scarce resource regime by setting *c*_1_ to 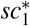, where *s* =1.01 or 1.05, implying that when resources are scarce, it takes longer (*i.e.*, more energy) for a group to detect each resource. For each combination of parameter values, we ran 10,000 simulations for each group size and computed the optimal group size for both the abundant and scarce resource conditions (see *Methods: Computing the optimal group size*). In our analyses, we only included a set of parameter values if at least one group size, in each of the abundant and scarce conditions, had a survival probability greater than 5%, and if none of the group sizes, in both the abundant and scarce conditions, had survival probabilities greater than 95%, in order to accurately estimate the optimal group size.

### Limits on the strength of the social benefits

We typically assume that the benefits of sociality increase sublinearly with group size. For the collective territoriality, collective niche expansion, and collective consumption mechanisms, we simply set 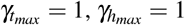, and 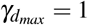. For other mortality, we need *p*_*p*_(*M* = 2) ≥ *p*_*p*_(*M* = 1)/2, so that 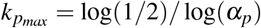.

We define the criteria for the sensing/detection, energetics, and dispersal mechanisms somewhat differently because the sublinear accrual of benefits cannot be defined for these mechanisms. Instead, for these mechanisms, we assume that groups of size 2 (*M* = 2) have, at best, a performance halfway between a solitary individual and an infinitely large group. For the sensing/detection mechanism, we have *p*_*s*_(*M* = 2) ≤ (1 + *α*_*s*_)/2, which gives us 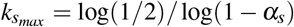. For the energetics mechanism, we have *c*_*b*_(*M* = 2) ≥ *c*_1_(1 + *α*_*b*_)/2, so that 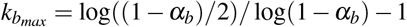. For the dispersal mechanism, we have *p*_*e*_(*M* = 2) ≤ *α*_*e*_/2, so that 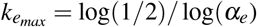.

### Calculating the “edge of starvation”

We define the “edge of starvation” as an environment where a solitary individual in the abundant resources condition has, on average, exactly 0 energy stores at the end of the simulation.

To calculate this, we first calculate the mean energy that a solitary individual extracts from a potential resource. The probability that a resource is of size *C* is 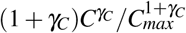, where we set *C*_*max*_ = 1 and *γ*_*C*_ = −3/4. There are two conditions to consider: 1) where *α*_*h*_ ≤ *α*_*d*_, and 2) where *α*_*h*_ > *α*_*d*_. For condition (1), a solitary individual can entirely consume any resource that it captures, so we just need to find the expected size of resources ranging in size from 0 to *α*_*h*_:

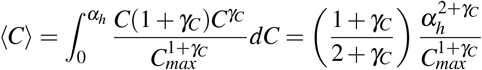

For condition (2), a solitary individual can entirely consume a resource it captures if its size *C* ≤ *α*_*d*_ but only consumes *α*_*d*_ of the resource if *C* > *α*_*d*_; the mean energy consumed in this case is:

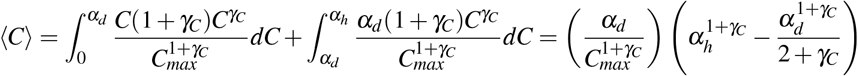

The mean total energy gained in a simulation by a solitary individual is *C*_*total*_ = *μ*_1_*α*_*s*_ < *C* >. For the solitary individual to have 0 energy at the end of the simulation, we need *c*_0_ + *C*_*total*_ −*c*_1_ = 0.

### Computing the optimal group size

For all of the mechanisms except the dispersal mechanism, we simply locate the group size, for each of the abundant and scarce resource conditions, that maximizes the probability of survival. For the dispersal mechanism, the probability of survival asymptotes to *α*_*e*_ for large group sizes, so it is difficult to determine the optimal group size in simulations because a wide range of group sizes have survival probabilities close to *α*_*e*_. We observe that, for this mechanism, as group size increases, the probability of survival appears to either monotonically increase, or initially decrease and then increase. Therefore, the optimal group size can only either be *M* = 1 or *M* = ∞ (or *M* = 100 for our simulations). Therefore, we simply observed the survival probability for the two extreme group sizes to determine the optimal group size.

### Analytical solution

Here, we assume *C*_*max*_ > *C*_*h*_. Let us use *P*_*c*_(*x*; *M*) as the probability that an individual consumes exactly *x* calories when in a group size *M*, and *F*_*c*_(*x*; *M*) as the probability function of *P*_*c*_(*x*; *M*)—*i.e.*, the probability that an individual consumes *x* calories or less (*F*_*c*_(*x*; *M*) = *P*_*c*_(*X* ≤ *x*; *M*)). Altogether, we may write the survival probability of an individual, *S*(*M*), as

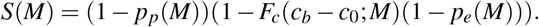

That is, an individual survives if it does not die from non-resource-related causes (with probability 1 − *p*_*p*_(*M*)) and it does not die of starvation, which we can calculate as the reciprocal of the probability of starvation—given by the probability that the caloric intake is insufficient (*F*_*c*_(*c*_*b*_ − *c*_0_; *M*))—times the probability of unsuccessful dispersal 1 − *p*_*e*_(*M*). As before, *c*_*b*_(*M*) denotes the energy that individuals burn and *c*_0_ = 1 represents the initial energy level—an individual has insufficient energy to survive when consuming less than *c*_*b*_ − *c*_0_. While *p*_*p*_(*M*) and *p*_*e*_(*M*) are directly set by model parameters (see *Methods: Detailed description of the model framework*), *F*_*c*_(*x*; *M*) is not; it is instead an emergent quantity of the model, which we now derive. To simplify the notation, we will omit arguments *M*, and simply refer to *p*_*p*_, *p*_*e*_, *c*_*b*_, *C*_*h*_, *c*_*d*_, and *F*_*c*_(*x*).

### *One prey scenario* (*μ*_1_ = 1 *and γ*_*t*_ = 0)

First, we note that an individual obtains no resources at all when 1) the available prey is larger than the niche size *C*_*h*_ or 2) even if its size is within reach, when the prey is not detected or captured. The probability of obtaining 0 resources is therefore given by

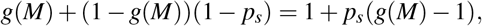

where *g*(*M*) is the probability that prey has size higher than *C*_*h*_, 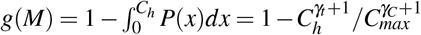 and *p*_*s*_ is the capture

Second, the probability of capturing prey, yet consuming *x* or less out of it, is given by 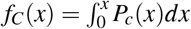, where *P*_*c*_(*x*) is the probability of consuming exactly *x* calories. Note that the prey captured is divided by the group (with size *M*) before being consumed. As a result, the probability of consuming *y* calories, *P*_*c*_(*y*), can be derived from the probability of capturing a prey worth *x* calories 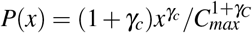, after the change of variable *y* = *x*/*M*, yielding *y*^−1^ = *h*(*x*) = *xM*. Accounting for the Jacobian involved in the change of variable, this results in *P*_*c*_(*x*) = *P*(*h*(*x*))*h’*(*x*) = *P*(*xM*)*M*. The probability of consuming less than *x* of a captured resource, *f*_*C*_(*x*; *M*), thus reads

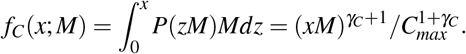

We can now write down the probability of consuming less than *x* calories, *F*_*c*_(*x*). It is given by 1) the probability of failing to capture any resource plus 2) the probability of capturing a resource yet consuming less than 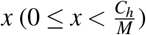 out of it:

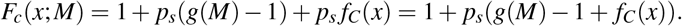

Finally, we complement the probability function above to account for consumption limitation due to the collective consumption mechanism. If *c*_*d*_ *M* > *C*_*h*_ (**scenario 1**), collective consumption does not play any role: it would only affect the level of energy intake for prey larger than the threshold imposed by the niche constraint. If *c*_*d*_ *M* < *C*_*h*_ (**scenario 2**) and *x* > *x*_*d*_, however, collective consumption—rather than resource competition—limits energy intake. This means that a prey can be so large that it cannot be fully consumed by the group. If this is the case, then an individual always consumes *c*_*d*_ or less, such that *F*_*c*_(*x*) is given by

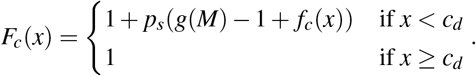

To summarize, the probability of survival is given by

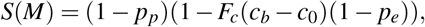

where under **scenario 1**(*c*_*d*_*M* > *C*_*h*_) the probability function of resource consumed is given by

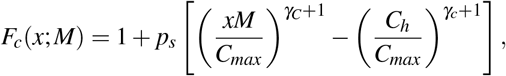

and, under **scenario 2**(*c*_*d*_ *M* < *C*_*h*_), the probability function of resource consumed is given by

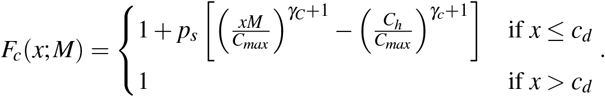

### Multiple prey scenario (*μ*_1_ > 1 *or γ*_*t*_ > 0)

When more than one resource is available, calculating *F*_*c*_(*x*) requires calculating the probability that consuming a combination of several resources leads to insufficient energy. Given the highly heterogeneous distribution of resource sizes assumed, such calculation involves a combinatorial problem hard to solve analytically. Nonetheless, it is possible to compute *F*_*c*_(*x*) numerically, when multiple resources exist. We can use the inverse transform sampling method to sample random values from a distribution of caloric gains per resource. Random values of caloric gain can be sampled by 1) sampling random values from a standard uniform distribution (*y* ~ *U* (0, 1)) and 2) calculating 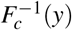, where 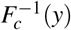 is the inverse of function *F*_*c*_(*x*) (*i.e.,* the CDF function of *P*_*c*_(*x*), the probability of consuming *x*—defined above). Considering the two scenarios already mentioned, we have, for **scenario 1**(*c*_*d*_ *M* > *C*_*h*_),

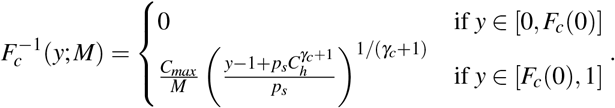

Note that *F*_*c*_(*x*) is only defined in the interval 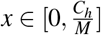 thereby 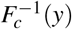 is defined for 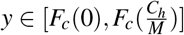, where 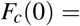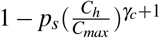 and 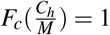. Following a similar reasoning, for **scenario 2**(*c*_*d*_ *M* < *C*_*h*_) we have:

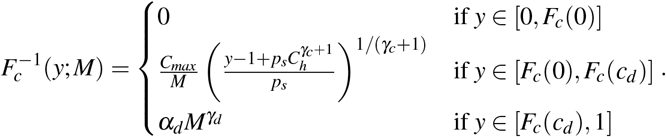

Where 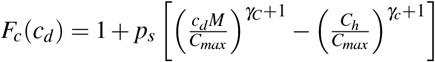. Using 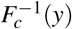, we can now calculate the probability of getting insufficient energy by taking the average (over many trials, *N*) of sampling *μ* values of energy intake, using 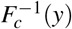 and each time sampling *y* from a uniform standard distribution. This way, a numerical approximation of *F*_*c*_(*x*; *M*)—which we denote by 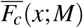—is given by

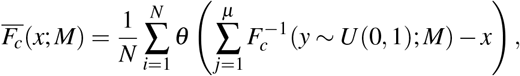

where *θ* (*x*) = 1 if *x* ≥ 0 and *θ* (*x*) = 0 otherwise. 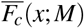 can be directly used in the probability of survival, *S*(*M*), defined above. We used *N* = 10, 000 in Figure 2A and *N* = 1, 000 in Figure 4.

### Dragon and unicorn analyses

Based on the supernatural histories of dragons and unicorns, we posited ranges for the parameters *μ*_1_, *α*_*s*_, *α*_*h*_, *α*_*d*_, *α*_*b*_, *α*_*p*_, and *α*_*e*_ for the two species. We assume that dragons eat large meals relatively infrequently, so we set *μ*_1_ = 1. When dragons do hunt we assume they detect prey relatively easily since they can fly, and that their fierceness allows them to capture even large prey solitarily—though not the very largest potential prey in their environment. We therefore restricted both the individual detection/capture parameter (*α*_*s*_) and the niche or maximum size parameter (*α*_*h*_) for dragons to between 0.5-0.7. Likewise, we also use a relatively high range of 0.5-0.7 for the individual consumption parameter (*α*_*d*_) for dragons, as their fire-breathing abilities mean that they can cook their food, to better extract nutrients and delay rot. We fixed *α*_*b*_ at 0.4, as dragons can take advantage to some degree of aerodynamic drafting while flying in groups, *α*_*p*_ at 0.3, as dragons’ extremely low predation risk reduces other mortality overall, and *α*_*e*_ at 0.8, as dragons’ flight makes them excellent dispersers. The last three parameters were fixed at specific values, rather than ranges, for both dragons and unicorns, so that all of the mechanisms took up the same volume of parameter space and could be directly compared with each other.

While one may expect unicorns’ powers to help them find resources, we assume their magical horn does not aid in foraging, specifically. We set *μ*_1_ = 1 for unicorns, as they typically will not encounter more than one patch of grass at a time. Unicorns are notoriously skittish, so we assume that they spend a large portion of their time on anti-predator vigilance, leaving less time for foraging. Therefore, we restrict *α*_*s*_ to between 0.1-0.3. We also restrict *α*_*h*_ and *α*_*d*_ to 0.1-0.3, as their herbivorous diet means that even their largest meals are relatively low in calories, and they can consume only a small fraction of a grass patch they find. We fixed *α*_*b*_ at 0.8, as unicorns derive no strong group benefit in terms of energetics, *α*_*p*_ at 0.5, because, while they are cryptic, they are susceptible to predators, and *α*_*e*_ at 0.2, as unicorns are known for rarely leaving the relative safety of their natal forest, thus making successful dispersal unlikely.

We used the analytical solution for this analysis, and uniformly sampled the region of parameter space for each species at a resolution of 0.01 for each parameter, recording whether the optimal group size decreased, increased, or did not change, when resources became scarce. For Figures 4C and F, we removed results for which the optimal group size was 1 for both conditions (because we are interested in the social behavior of these two species). Then, given a particular hypothetical experimental result (*i.e.*, decrease, increase, or no change), we calculated the proportion of results arising from each of the seven benefits of sociality.

